# PTM-Driven Reshaping of the Peptide Translocation Landscape in Bilayer Graphene Nanopores

**DOI:** 10.64898/2026.01.26.701721

**Authors:** Anurag Upadhyaya, Pranjal Sur, Manoj Varma, Prabal K. Maiti

## Abstract

Post-translational modifications (PTMs) underpin much of protein regulation, yet their single-molecule readout remains a challenge in nanopore proteomics. While biological nanopores have shown exquisite PTM sensitivity, the microscopic mechanisms by which PTMs perturb signals in solid-state nanopores are largely unexplored. Here, we use all-atom molecular dynamics to investigate how three common PTMs, acetylation, phosphorylation, and methylation, modulate the translocation of a cancer-relevant p53 peptide fragment through a bilayer graphene nanopore. We find that PTMs remodel the translocation landscape far more strongly at the level of dwell-time statistics than at the level of mean current blockade. Acetylation enhances peptide–graphene adhesion and substantially slows transport, with adjacent acetylations producing the longest residence times due to cooperative interfacial interactions, while remotely spaced acetylations yield broader, heterogeneous dynamics. Phosphorylation introduces a negative charge that increases dwell time through an electrostatic tug-of-war, while also generating the largest current blockade among the PTMs studied. In contrast, methylation minimally perturbs translocation due to weak pore interactions and preserved charge. Combining dwell time with relative blockade features enables a simple linear SVM classifier to reliably distinguish unmodified, acetylated, and phosphorylated states. These results establish mechanistic design principles for PTM detection using solid-state nanopores and delineate which classes of PTMs are the most amenable to single-molecule detection with these devices.

## Introduction

Post-translational modifications (PTMs) are critical biochemical alterations experienced by proteins after translation, profoundly influencing their function, localization, stability, and interaction with other molecules. These modifications, which include acetylation, phosphorylation, ubiquitination, glycosylation, and methylation among others, play vital roles in regulating cellular processes and maintaining homeostasis^[1–5]^. Identifying PTMs is essential for understanding protein function and for the development of therapeutic interventions, as aberrant PTMs are often associated with diseases such as cancer, neurodegenerative disorders, and metabolic syndromes^[1,5–8]^. Mass spectrometry is one of the conventional methods available for detecting PTMs^[9,10]^. However, detecting PTMs from mass spectrometry data is challenging due to the proteome’s complexity, wide range of protein abundance, chemical diversity of modifications and the requirement of substantial amount of sample inputs. These limitations often hinder mass spectrometry’s ability to precisely detect the positions of PTMs, especially among multiple sites.

Nanopore detection of PTMs is an emerging area of research that leverages nanopore technology for the identification and characterization of PTMs at the single-molecule level^[11– 21]^. Nanopore technology utilizes nanometer-sized pores through which individual biomolecules, such as DNA, RNA, and proteins, can pass. As molecules traverse through the nanopore, they cause characteristic changes in ionic currents, which can be detected and analyzed to infer various molecular properties including size, shape, charge, structure, conformation, orientation and modifications^[22–41]^. One intriguing aspect of PTMs is their impact on the translocation dynamics of peptide sequences through nanopores. Distinguishing the structural differences between modified amino acids using solid state nanopores is challenging due to their high translocation speed and small volume differences introduced by modifications^[42]^. The relative dearth of studies, especially theoretical ones, focusing on the detection of PTMs using solid-state nanopores motivates our study. Specifically, we seek to understand how three major PTMs, namely, phosphorylation, acetylation, and methylation impact peptide translocation through a bi-layer graphene nanopore. Previous literature has shown that the flicker noise produced by a graphene nanopore tends to reduce with increasing layer numbers^[43]^. Moreover, with increasing thickness, higher dwell time has been reported^[44]^, motivating us to study nanopores on bilayer graphene membrane.

We chose a peptide segment from p-53, a biologically crucial protein for cell cycle regulation, as our analyte of interest. The sequence containing residues 365 to 375 in C-terminus of p-53, is subjected to various PTMs such as phosphorylation, acetylation, and methylation, which can modulate the activity, stability, and interactions of the p53 protein^[45– 55]^

We used all-atom molecular dynamics (MD) simulations to probe the trajectory of the peptides, to extract the dwell time of the peptide inside the pore and to calculate the ionic current blockade during translocation. The PTMs we considered are the following: (a) single phosphorylation (Ph) at SER (367), (b) single methylation (Met) on LYS, (c) double methylation (Met_double) on LYS (370,372), (d) single acetylation (Ac) on LYS (370), (e) double acetylation apart (Ac_apart) on LYS (370,373), where the acetylation cites are remotely spaced, (f) double acetylation adjacent (Ac_adjacent) on LYS (372,373), where the acetylation cites are closely spaced. The choice of these PTMs allows us to study the effect of the number as well as position of the PTM as this is known to be an important factor in human diseases^[56]^related to protein disfunctions. For example, activation of sequence specific DNA binding of p53 is significantly dependent on synergistic acetylation^[57]^ on specific positions in its C-terminal and low number of these acetylations can be linked with cancer. Hence, the double modifications that differ by positions were introduced as we wanted to investigate if our method is sensitive enough to detect the numbers as well as the positions of PTMs. From these studies, we found that different PTMs alter graphene-peptide interactions, leading to distinct translocation dynamics resulting in distinct dwell time variations. Based on the dwell time estimates, we can identify phosphorylation, methylation, and closely and remotely spaced acetylation. Although, different PTMs produce distinguishable relative current blockade values, the different dwell times for various PTMs appear to be more significant. Moreover, to test the robustness of the dwell time-based detection, we performed SVM^[58]^ based classification among different PTMs. Based on the dwell times and relative current blockades, we could distinguish phosphorylation, methylation, and acetylation, including whether it was closely or remotely spaced with reasonable accuracy using the SVM classifier. Our work demonstrates that different PTMs alter graphene-peptide interactions, leading to distinct translocation dynamics and consequently resulting in distinct dwell time variations. **Fig.1**(a, c) illustrates the peptide segment used, including the residues and their possible post-translational modifications. We attached a homopeptide of five arginine residues (ANCHOR) to both the ends of our peptide segment from p-53. The role of the positively charged ANCHORs was to facilitate the translocation of the p-53 segment.

**Figure 1:**
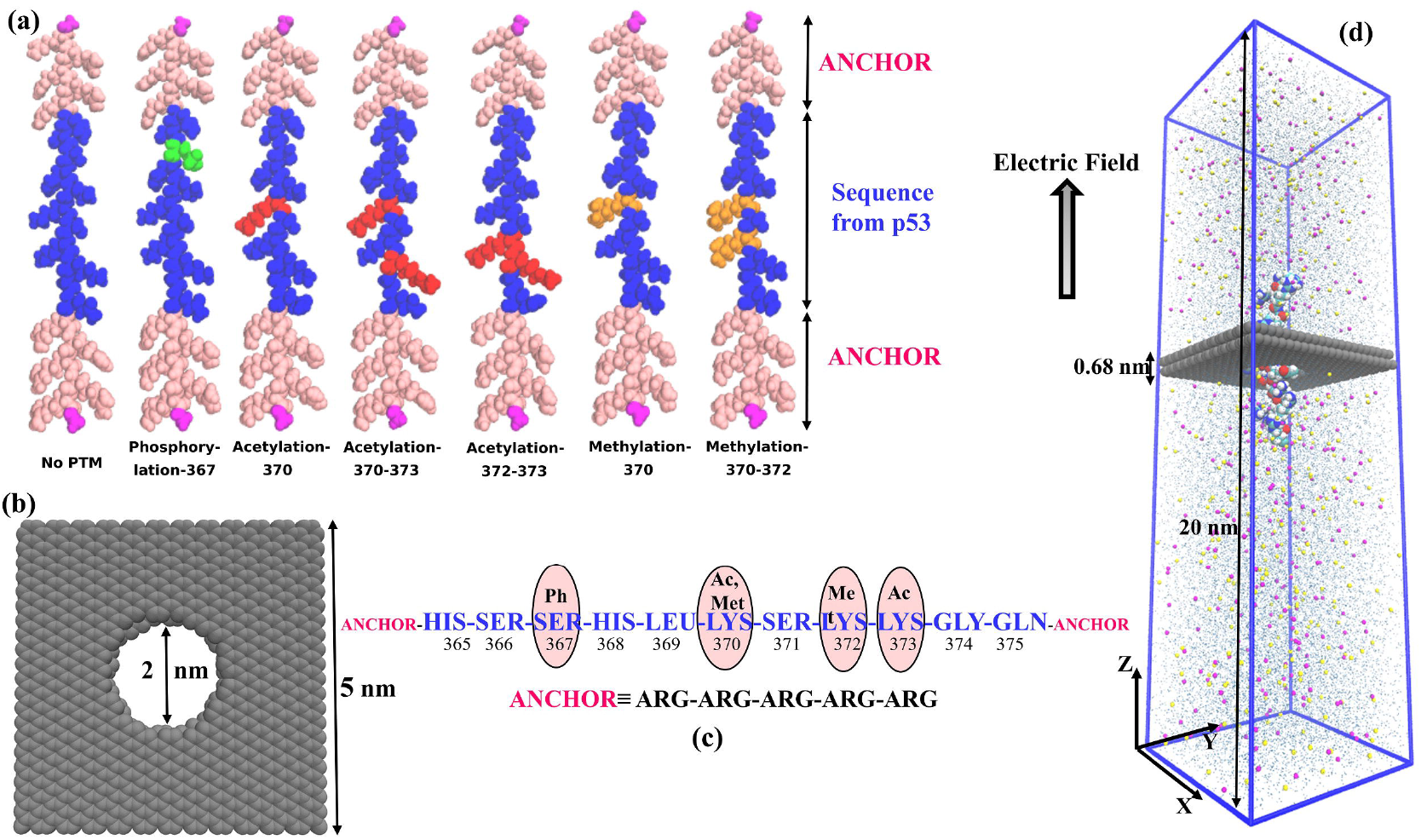
**(a)** The design of the peptides subjected to translocation. The blue residues are the segments taken from C-terminus of p53. The pink residues are positively charged ARG anchors. The role of the anchors is to help the p53 segment translocate through the pore. The magenta residues are NME and ACE capping in the terminal residues in the anchor. The different PTMs have been colored differently. **(b)** The graphene bilayer membrane with a pore of 2nm diameter, in its center. **(c)** Sequence of the C-terminus of the p53 protein from residue number 365 to 375. The PTM sites have been circled. **(d)** The schematic of the set-up of our simulation. A peptide is translocating through the graphene nanopore.

## Results and Discussions

### Dwell time as a dominant discriminator of PTMs

We compare the calculated dwell time and relative current blockade for peptides subjected to different PTMs in **Fig.2a** and **Fig.2b** respectively. Our data demonstrates that dwell time exhibits greater sensitivity than relative current blockade. The dwell time histograms have been fitted with lognormal distribution to obtain the probability density function shown in **Fig. 3a**. The pronounced right skewness of the dwell time distribution indicates heterogeneity in translocation dynamics, reflecting the absence of a single characteristic timescale. This suggests that the underlying kinetics are governed by stochastic pore–analyte interactions, energy barrier crossings, conformational heterogeneity of the analyte etc. In contrast, the relative current blockade histograms are well described by a normal distribution (**Fig. 3b**), implying that the decrease in current is rather homogeneous primarily influenced by additive noise. The goodness of the fits has been discussed in SI (section 2, **Table S4)**.

**Fig. 2.**
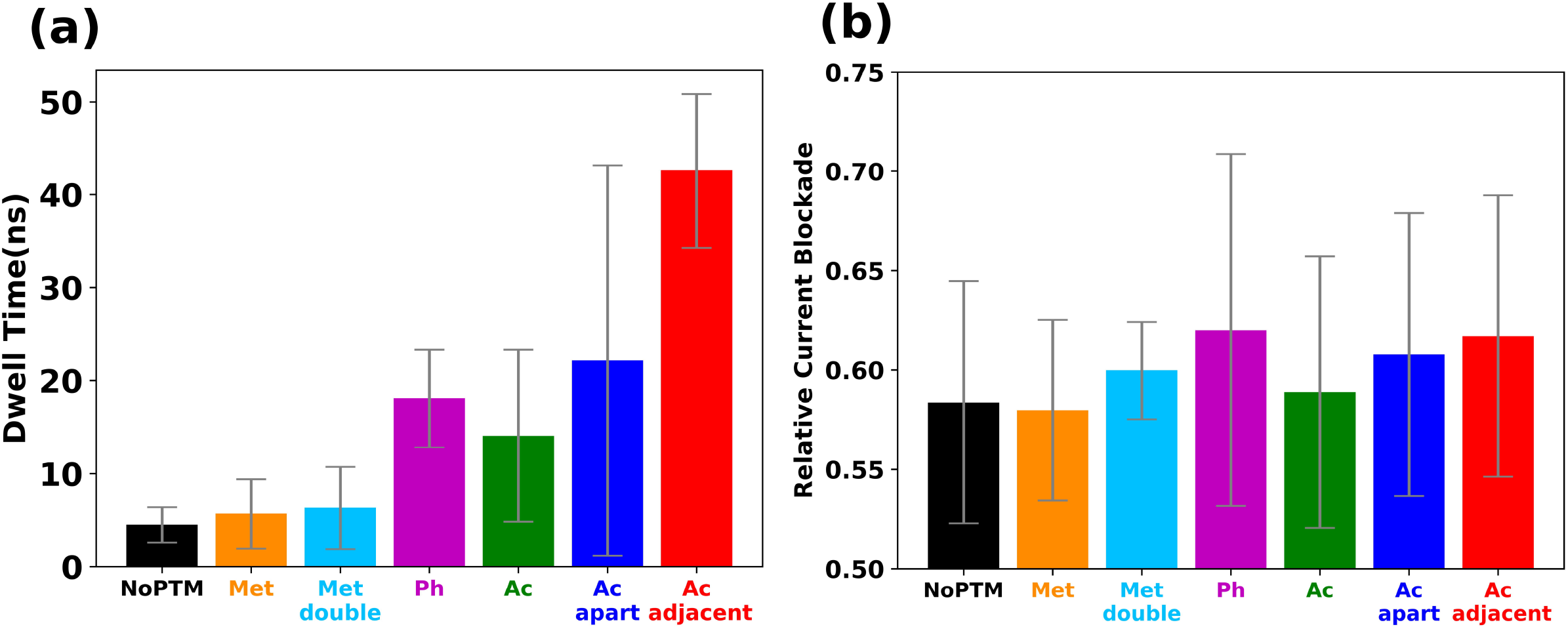
**(a)** Bar plot demonstrating difference in average dwell time for differently modified peptides. Prominent distinction is present among methylation, phosphorylation, and acetylation. In addition, three variations of acetylation have distinct differences in their dwell times. Double acetylations have greater dwell time than single acetylations. Furthermore, closely spaced acetylation exhibit prolonged residence compared to remotely spaced acetylation. **(b)** Bar plot demonstrating difference in average relative current blockade for differently modified peptides. Phosphorylation, which introduces a net negative charge in modification site, exhibit the highest relative current blockade even though its residence at the pore is not the highest, possibly highlighting the importance of electrostatic interaction and local charge distribution in governing current blockades. The current blockade increases as the number of PTM increases since double methylation and acetylation exhibit higher relative current blockade compared to single methylation and acetylation, respectively.

**Fig. 3.**
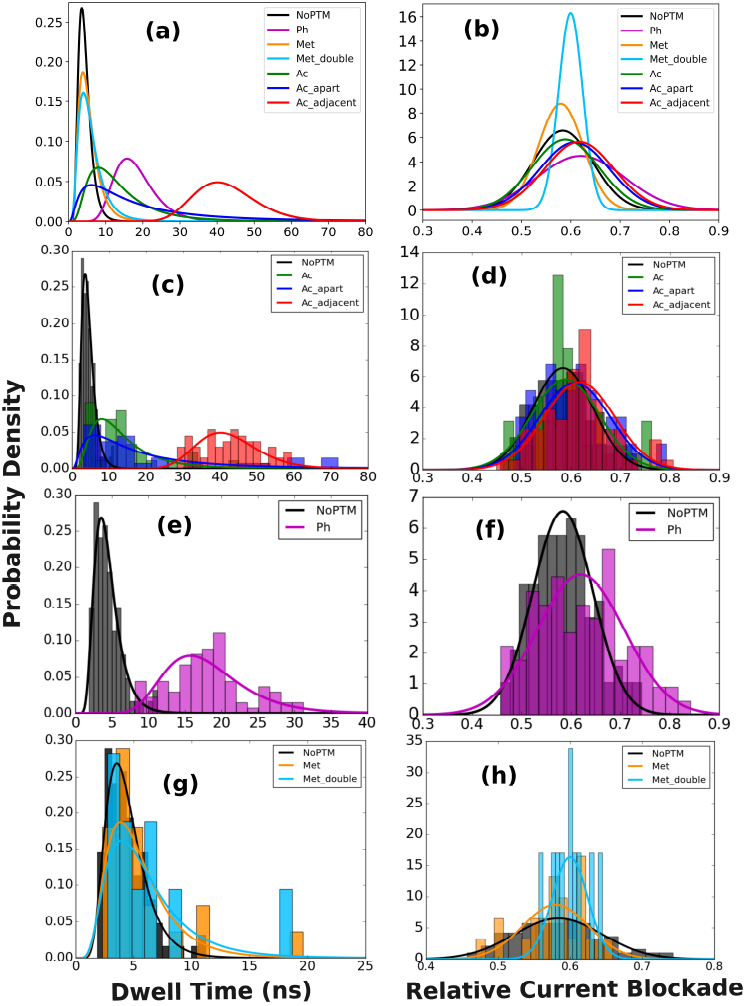
**(a)** The probability density distribution of dwell time in case of different PTMs. **(b)** The probability density distribution of relative current blockades subjected to different PTMs. **(c)** The comparison of histograms of dwell time with the fitted probability density for NoPTM and all the acetylation (Ac) variants. The plots are well distinct for each variant. The NoPTM shows least spread while the acetylated peptides exhibit variety in translocation dynamics. **(d)** The comparison amongst histograms of relative current blockade with the fitted probability density for NoPTM and all the acetylation variants. **(e)** The comparison between histograms of dwell time, **(f)** relative current blockade with the fitted probability distribution for phosphorylation (Ph) and NoPTM. Both dwell time and relative current blockade for Ph is prominently larger. **(g)** The comparison amongst histograms of dwell time, **(h)** relative current blockade with the fitted probability distribution for Met, Met_double, NoPTM. The dwell time does not differ much, Met_double exhibiting the highest value marginally. The histograms are fitted with lognormal distribution for dwell time and normal distribution for current blockades.

The major effects of the different PTMs on peptide translocation are discussed below.

### Effect of acetylation on translocation: mechanistic origin of long dwell times

Acetylation (Ac) of LYS residue neutralizes the +1e charge of LYS. Since positive charge helps the whole peptide to move in the direction of the electric field, neutralization of LYS contributes to diffusive dynamics and reduces field driven dynamics. Moreover, acetylated residues exhibit a strong interaction with graphene surface. Hence, more the number of acetylation, greater is the dwell time of the peptide as evident in **Fig.2a, Fig.3a, c**. These figures also tell us that, not only the dwell time of double-acetylated (Ac_apart, Ac_adjacent) peptides is greater than mono-acetylated (Ac) ones, even within the double-acetylated PTMs, there is a distinct difference in dwell time depending on the positions of the two acetylations. **Fig.2b, Fig.3b, d** shows that relative current blockade is increased in the case of double acetylation compared to single acetylation, which agrees with the data shown in a previous study ^[13]^ using biological nanopores.

### Effect of phosphorylation on translocation: role of electrostatics

Phosphorylation (Ph) of neutral SER residue introduces a net −1e charge which opposes the drift of the peptide towards electric field, and a tug-of-war can be observed around pore during translocation while the phosphorylated residue itself avoids interaction with graphene (**Video S1**). The electrostatic repulsion opposes the drift of the peptide and enhances the dwell time. While Ph does not exhibit the highest dwell time (**Fig.2a**), it produces the largest average relative current blockade (**Fig.2b**). This highlights the role of local charge distribution and electrostatic interactions in governing the current blockade magnitude. Moreover, this underscores that dwell time and current blockade encode complementary physical information. **Fig.3e, f** demonstrates the histograms of dwell time and relative current blockade respectively, showing the distinction among NoPTM and Ph.

### Effect of methylation on translocation: minimal perturbation

**Fig.3g, h** shows the histograms of dwell time and relative current blockade for Methylation (Met), Met_double and NoPTM, exhibiting minimal distinction (**Fig.S3**) among NoPTM and methylation variants. Methylation of LYS residues do not introduce any additional charge, neither do they neutralize +1e charge of LYS. Since, the peptides are translocating in the direction of electric field, methylation does not resist the field driven movement of the peptide. Moreover, methylated LYS residues minimally interact with pore or membrane surface (**Video S2, S3**). Hence, the translocation dynamics of NoPTM and methylated peptides are similar, and we observe similar dwell time for NoPTM, Met and Met_double (**Fig.2a; Fig.3g, h**). In addition, we notice that the relative current blockade increases as the no. of methylation increases (**Fig.2b**). Although, the dwell time of methylated and acetylated residues are in stark contrast, their relative current blockade magnitude is comparable **(Fig.2a, b)**.

### Residue-level translocation dynamics and classification of PTM states

Next, we looked at residue-by-residue translocation of these peptides by tracking the position of the C_α_ atom of each residue inside the pore width. As we see from **Fig. 4**, acetylation (Ac) and phosphorylation (Ph) causes substantial slowing down of the residues in the neighbourhood of the PTM. We note that such slowing down is absent in the case of methylation (Met). These figures provide microscopic visualizations that suggests the origin of the dwell time distributions shown in **Fig. 2** and Fig. **3**. Trajectories corresponding to each case in **Fig.4** can be visualized in supplementary Video **S1, S3, S4, S10**.

**Fig. 4.**
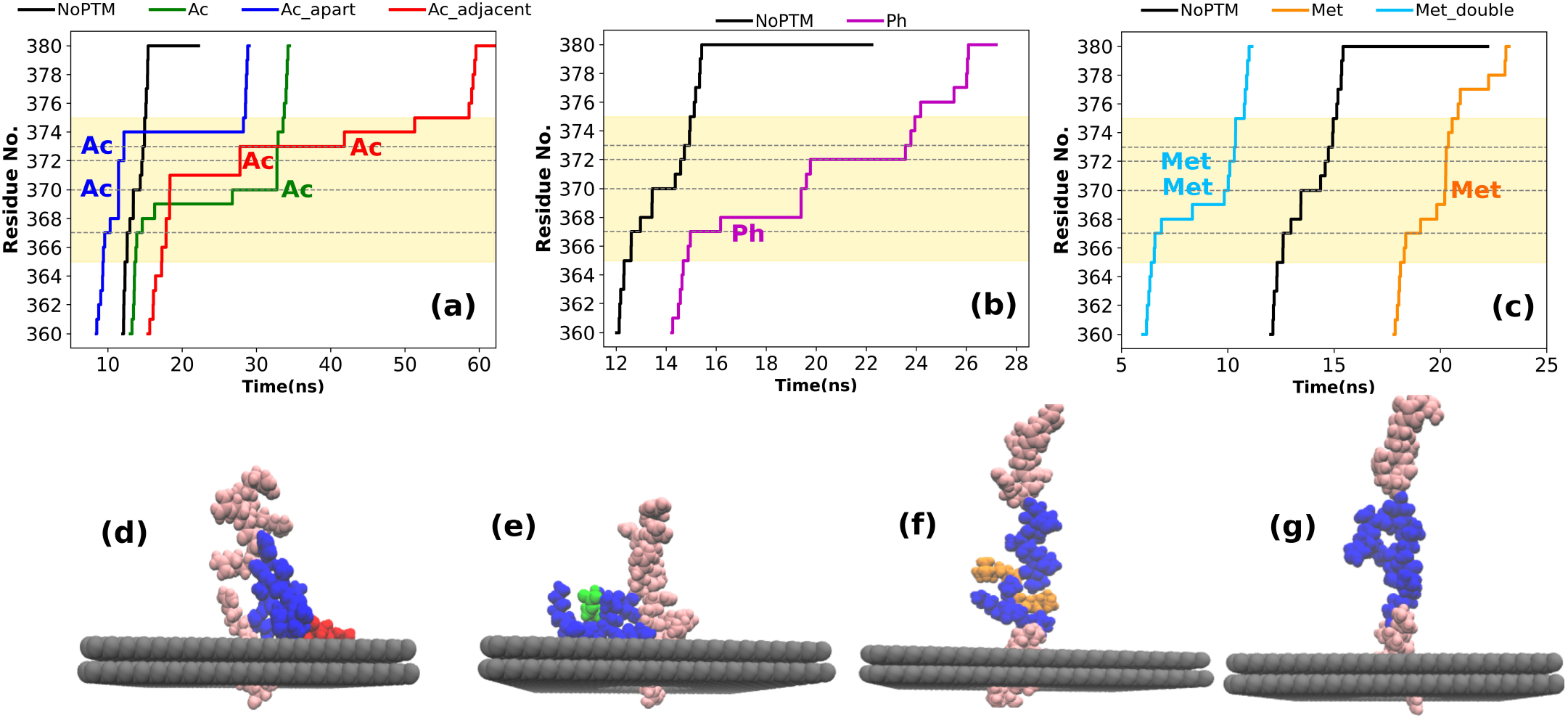
The yellowish shaded region is the translocation regime of the p-53 sequence and the dwell time of translocation is the difference of start and end of this region in time axis. **(a)** Residue-wise translocation of Ac, Ac_apart, Ac_adjacent along with NoPTM. Larger translocation steps in time axis are observed at the acetylation site or their immediate neighboring sites compared to when the anchors or the non-neighboring residues of acetylation site are translocating. The dwell time of Ac_adjacent is significantly higher compared to the other cases. **(b)** Residue-wise translocation, for phosphorylation and NoPTM, **(c)** for Met, Met_double NoPTM. No significant increase in translocation steps about the methylated sites as compared to acetylated and phosphorylated ones **(d)** End stage of translocation for peptides subjected to different PTMs **Ac**: The acetylated residue adheres to the membrane. Translocation kinetics dominated by pore-analyte interaction **(e) Ph**: Although, the phosphorylated residue avoids direct interaction with membrane (**video S1**), the overall coiling over membrane indicates existence of surface-peptide interaction while translocating. Translocation kinetics dominated by interplay between electrostatic attraction-repulsion. **(f) Met_double:** The methylated residues do not interact with pore/membrane significantly. The peptide is rather stretched than coiled at the end stage of translocation. **(g) NoPTM:** The peptide with no PTM, exhibit minimal pore/surface interaction. Both Met_double and NoPTM exhibit field driven translocation dynamics.

We found that a simple classifier such as Support Vector Machines (SVM) can distinguish the differences between the various PTMs based on their translocation behavior, specifically the dwell time and the relative blockade current. **Fig.5a** shows that the classification among NoPTM and different variants of acetylations can be performed with an overall accuracy of 84.8%. The loss of accuracy is primarily caused by the poor identification capability for Ac_apart. We observe that our model correctly identified Ac, Ac_adjacent with a 100% accuracy, detected the unmodified sequence with 93.3% accuracy with a 6.7% confusion with Ac. But accuracy for Ac_apart is 0%. The model classifies Ac_apart as either a NoPTM or an Ac_adjacent, with an equal confusion of 50%. **Fig.5b** tells us that Ac_apart does not have a well-defined decision space and our model fails to identify Ac_apart as a separate class. On other hand, the NoPTM, Ph and Ac_adjacent can be distinguished with an overall accuracy of 94%. (**Fig.5c**). They each have a distinct domain in relative current blockade-dwell time decision space (**Fig.5d**).

**Fig. 5.**
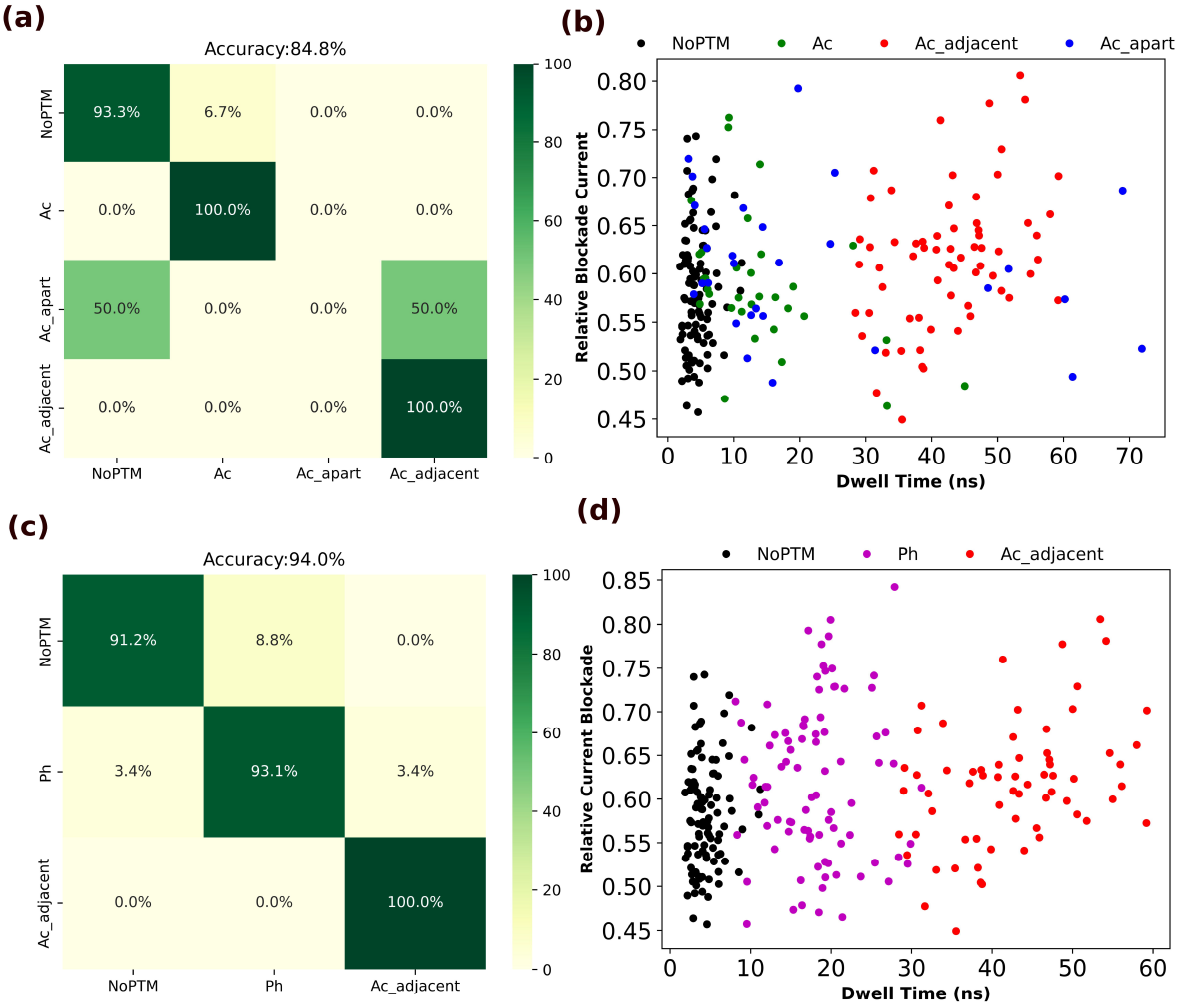
**(a)** Confusion matrix calculated using SVM, subjected to classification based on dwell time and relative current blockade. NoPTM and three types of acetylation are classified here. Classification accuracy of NoPTM, Ac, Ac_adjacent is reasonably well. Although, Ac_apart is being classified as Ac or Ac_adjacent. **(b)** Relative current blockade vs dwell time for NoPTM and three types of acetylation. We can see the spread of the blue points, corresponding to Ac_apart, which caused the classification confusion in (a). **(c)** Confusion matrix for a SVM classification among NoPTM, Ph and Ac_adjacent. All the three classes are well distinct. **(d)** Relative current blockade vs dwell time for NoPTM, Ph and Ac_adjacent.

### Positional heterogeneity and classification limits for acetylated peptides

For a better understanding of the position dependence of acetylation on the dwell-time, we inspected the individual trajectories. Acetylations that are adjacent or closely spaced tend to adhere to the graphene pore or membrane almost synchronously, resulting in strong interfacial interactions (**Fig. 6c, e**). In contrast, when acetylations are separated by two residues, their adhesion behavior becomes more heterogeneous. At times, their dynamics are dominated by drift–diffusion due to weak interactions between the acetylated residues and graphene (**Fig. 6d**). The dwell time for these types is in lower range for Ac_apart. Occasionally, their translocation dynamics are strongly governed by membrane adherence (**Fig. 6f**), leading to high dwell time similar to that observed for adjacent acetylations. In other instances, one acetylated residue adheres to the membrane while the other does not (**Fig. 6h**), leading to intermediate translocation time between the two extremes. Overall, Ac_apart displays increased heterogeneity in translocation pathways relative to other acetylation variants. This explains why we observe a spread in **Fig. 5b** and our model is not able to identify Ac_apart as a distinct class.

**Fig. 6.**
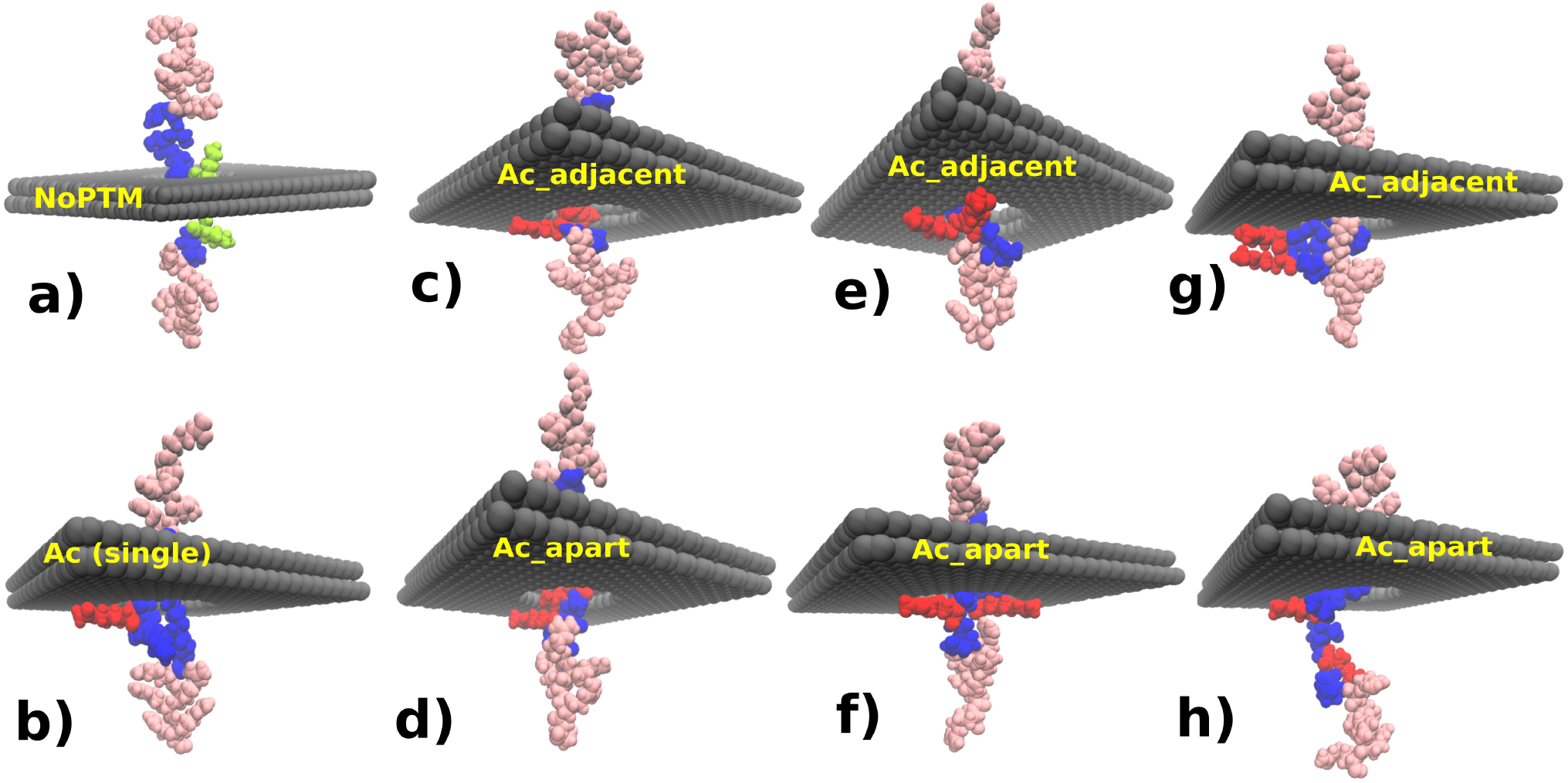
Variation in pore-peptide interaction among NoPTM and three different types of acetylations. **(a)** In NoPTM, the non-acetylated Lys residues (green) do not adhere to the graphene surface. Pore-peptide interaction is not significant. **(b)** Single acetylated residue (red) simply gets adhered to the graphene membrane while translocating, increasing dwell time compared to NoPTM. **(c–d)** In Ac_adjacent and Ac_apart, the two acetylated residues do not always bind simultaneously: one may align at the pore mouth while the other adheres outside. Although the conformations appear similar, Ac_adjacent exhibits greater dwell times, highlighting the role of residue proximity. **(e-f)** Cases where both acetylations bind the membrane together produce strong and long, comparable dwell times for Ac_adjacent and Ac_apart. **(g-h)** These conformations show only one acetylated residue binding, where the first acetylated residue that comes to pore mouth gets adhered to the membrane like **b** and the following one doesn’t. In Ac_adjacent, stacking of the two residues can keep one persistently attached, increasing dwell time, whereas in Ac_apart the unbound residue is more mobile, and the dwell time is somewhat between **d** and **f**. Overall, Ac_apart yields broader dwell-time variability compared to Ac_adjacent. Neighbouring acetylations enhance the dwell time despite conformational variances. Examples are provided in Video S4-S9.

## Conclusions

We present a systematic molecular-level analysis of how common PTMs influence peptide transport through graphene nanopores. Our results establish dwell time as the most sensitive and physically interpretable metric for PTM detection, surpassing relative current blockade in discriminative capability. Acetylation and phosphorylation perturb translocation strongly but through distinct mechanisms: acetylation enhances membrane adhesion and slows transport through interfacial interactions, whereas phosphorylation modulates dynamics through charge-driven electrostatic forces while generating the largest current signatures. Methylation, by contrast, induces only minor perturbations, underscoring that not all PTMs provide strong solid-state sensing contrast. Crucially, the positional arrangement of PTMs, illustrated by adjacent versus remotely spaced acetylations, produces clear, quantifiable differences in dwell-time distributions. This highlights that solid-state nanopores possess sufficient spatial sensitivity to report on both the number and position of chemically similar PTMs, a feature directly relevant to disease-linked PTM patterns in proteins such as p53. Finally, simple machine-learning classifiers can leverage simulation-derived features to distinguish multiple PTM states with high accuracy, demonstrating the robustness of the underlying physical signatures. Together, these findings provide mechanistic insights and practical design guidelines for developing next-generation solid-state nanopore sensors capable of resolving PTMs at the single-molecule level. They also identify which PTM classes are most amenable to detection using solid-state nanopores. This information will be valuable for experimental nanopore engineering and for expanding the scope of solid-state nanopore based protein sensing and sequencing technologies.

## Materials and Methods

We performed atomistic MD simulations using GROMACS^[59]^. CHARMM36m^[60]^ force field was used to describe all inter-and-intra-molecular interactions.

The total interaction energy can be written as sum of bonded and non-bonded interaction.

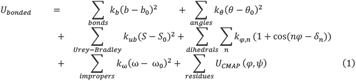

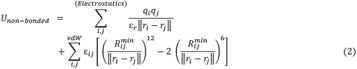

The parameters *b, θ, S, φ* and ω represent bond lengths, valence angles, Urey-Bradley 1,3-distances, dihedral angles, and improper torsion angles, respectively. The corresponding equilibrium values are denoted as *b*_0_, *θ*_0_, *S*_0_, ω_0_, while *k*_*b*_, *k*_*θ*_, *k*_*ub*_, *k*_*φ*_ represent the associated force constants. For the dihedral term, *k*_*φ*_,*n* is the amplitude, *n* is the multiplicity, and *δ*_*n*_is the phase shift. Additionally, U_CMAP_ accounts for backbone torsion corrections. Non-bonded interactions were calculated using partial atomic charges q_i,_ q_j_ with a dielectric constant of *ε*_*r*_. The van der Waals interaction between atoms i and j was modeled using the Lennard-Jones potential, where *ε*_*ij*_ is the effective potential well depth and 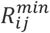 is the effective equilibrium distance. These effective parameters were derived using Lorentz-Berthelot combination rules, where ε_*ij*_ is the geometric mean of ε_*i*_ and ε_*j*,_ and 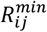 is the arithmetic mean of 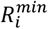 and 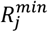. The term ‖*r*_*i*_ − *r*_*j*_‖ represents the Euclidean distance between atoms.

A pore diameter of 2 nm was created in the graphene bilayer (AB stacked) sheet with an interlayer spacing of 6.8Å. The bilayer graphene was position restrained throughout the simulations. The whole analyte including the p53 segment and the anchor was prepared using xleap module of AMBER^[61]^.

The simulation box dimensions were 5.3 x 5.3 x 20 nm. The bilayer nanopore was positioned at the center of the XY plane located at the mid-height of the box, while the peptide along with the anchor was placed below the pore in a way that the center of mass of the nearest anchor residue to the pore is 2.1 nm away from the pore. The rectangular simulation box containing the nanopore and peptide, was solvated with water. K^+^ and Cl^-^ ions were added to create a 1M KCl solution. Details of the system and runs can be found in SI (**Table S1, S2**). Periodic boundary conditions were applied in all three directions. We used the CHARMM modified TIP3P^[62,63]^ water model. The Settle^[64]^ algorithm was utilized to maintain the rigidity of the water molecules along with LINCS^[65]^ for inclusion of hydrogen bond constraints. CA atom type was assigned to model the graphene carbon atoms (charge neutral membrane). The interaction parameters for ions were those developed by Beglov and Roux^[66]^.

Energy minimization of the system was performed using the steepest descent method for 50,000 steps. The minimized system was then equilibrated at 300 K for 2 ns in the NVT ensemble using the Bussi-Donadio-Parinello thermostat^[67]^ (time constant=0.1ps). Subsequently, NPT equilibration was conducted for 2 ns to achieve equilibrium density at 1 bar pressure using an isotropic Berendsen^[68]^ barostat (time constant=2ps). Long-range electrostatic interactions were calculated using the Particle Mesh Ewald (PME) ^[69]^ summation technique. A cut-off distance of 12 Å was used for computing short-range Lennard-Jones interactions and the short-range part of Coulomb interactions. The production simulation was performed in the NVT ensemble using the Bussi-Donadio-Parinello thermostat, until the peptide is completely translocated, as described in the next section. The system temperature was maintained at 300 K. The integration time steps were 2 fs during both equilibration and production phases, and the trajectory was recorded at 2 ps intervals for analysis. An electric field of 0.05 V/nm was applied along the +Z direction in each simulation during the production run, creating a total of 1V bias across 20nm long simulation box. A restraint of 1000 kJ/mol.nm^2^ was applied to the C_α_ atom of every anchor residue (**Fig.1**(a, c)) during the production run. The PTMs were added to the peptide using CHARMM-GUI^[70]^. For phosphorylation, acetylation, and methylation, SP1, ALY and M3L residue type from Charmm36m has been used respectively. SP1 is the monophosphorylated form of SER, ALY is the acetylated form of LYS and M3L is the methylated form of LYS (details in **Table S3**). The dwell time was defined as the duration between the initial entry (T=t_1_) of the first residue of the sequence from p53 into the pore and the subsequent exit (T=t_2_) of the last residue of the sequence. The time-dependent ionic current^[24,71]^ through the nanopore was calculated using,

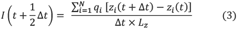

Where *I* is the ionic current, Δ*t* is 2 ps, and z_*i*_ is the z coordinate of the *i*^*th*^ ion, and *q*_*i*_ is its charge. N is the total number of ions, and the sum is for all the ions. Initially, the center of mass (COM) of the whole peptide was placed 6nm below the pore. When the electric field is switched on, the whole peptide starts moving towards the pore and eventually the translocation starts (**Fig.1d**). We calculated the blockade current, which is the average of the current (*I*_*avg*_)in the dwell time window. The relative current blockade has been defined as (*I*_0_ − *I*_*avg*_)/*I*_0_, where *I*_0_ is the open-pore current. In **Fig.S1, S2**, we show a typical current trace from our simulation. We used the Visual Molecular Dynamics (VMD)^[72–74]^ package to visualize the trajectories. For processing the trajectory and calculating dwell time and relative current blockade, we utilized MDAnalysis ^[75,76]^.

## Data, Materials and Software Availability

The data that support the findings of this study are available in the manuscript and supplementary information. Trajectories are available from the corresponding authors upon reasonable request.

## ACKNOWLEDGEMENTS

This work was supported by the Scientific and Useful Profound Research Advancement (SUPRA) Program of the Science Engineering Research Board (SERB) under Grant SPR/2021/000275 and the Scheme for Translational and Advanced Research in Science (STARS) under Grant MoE-STARS/STARS-2/2023-0077. P. S. acknowledges the PhD fellowship support from DST, India. We thank Prof. Anand Srivastava and Prof. Ashok Sekhar from Molecular Biophysics Unit, Indian Institute of Science, for insightful discussions.

## Notes

### Competing Interest Statement

The authors have declared no competing interest.

