## Supplementary Information for "PTM-Driven Reshaping of the Peptide Translocation Landscape in Bilayer Graphene Nanopores"

**Table S1: Number of atoms in the System**

| **K^+^** | **Cl^-^** | **Water** | **Graphene** | **Peptide** | **Total** |
| --- | --- | --- | --- | --- | --- |
| 338 | 349-351 | 50634-50691 | 1774 | 428-446 | 53525-53600 |

**Table S2: Number of production replicas for each PTM variant**

| **Ac** | **Ac_adjacent** | **Ac_apart** | **Ph** | **Met** | **Met_double** | **NoPTM** |
| --- | --- | --- | --- | --- | --- | --- |
| 32 | 65 | 29 | 88 | 24 | 10 | 100 |

**Table S3: PTM details**

| **PTM Type** | **Residue Type in Charmm36m** | **Modification on side chain** | **Charge introduced** | **Net charge of modified residue** |
| --- | --- | --- | --- | --- |
| **Acetylation** on LYS | ALY | LYS:  -(CH_2_)_4_-NH_3_^+^  ALY:  -(CH_2_)_4_-NH-C(=O)-CH_3_ | -1e | 0 (altered) |
| **Methylation** on LYS | M3L | LYS:  -(CH_2_)_4_-NH_3_^+^  M3L: -  -(CH_2_)_4_-N^+^(CH_3_)_3_ | 0 | +1e (unaltered) |
| **Phosphorylation** on SER | SP1 | SER:  -CH_2_-OH  SP1:  -CH_2_-O-P(=O)(OH)(O^-^) | -1e | -1e (altered) |

**1. Current Trace**

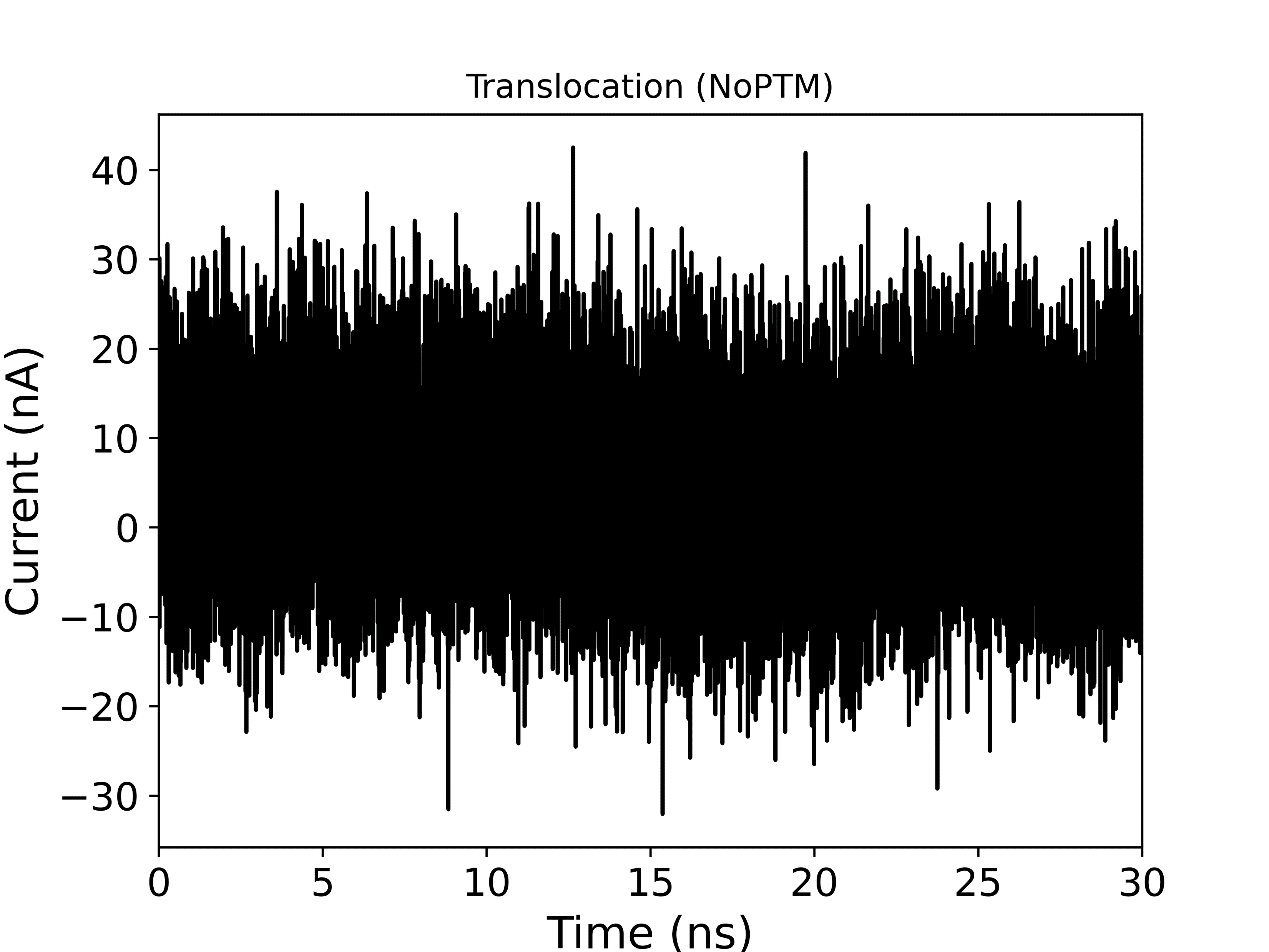

**Fig.S1**: Raw current signal generated using each frame of production run during translocation of a NoPTM peptide.

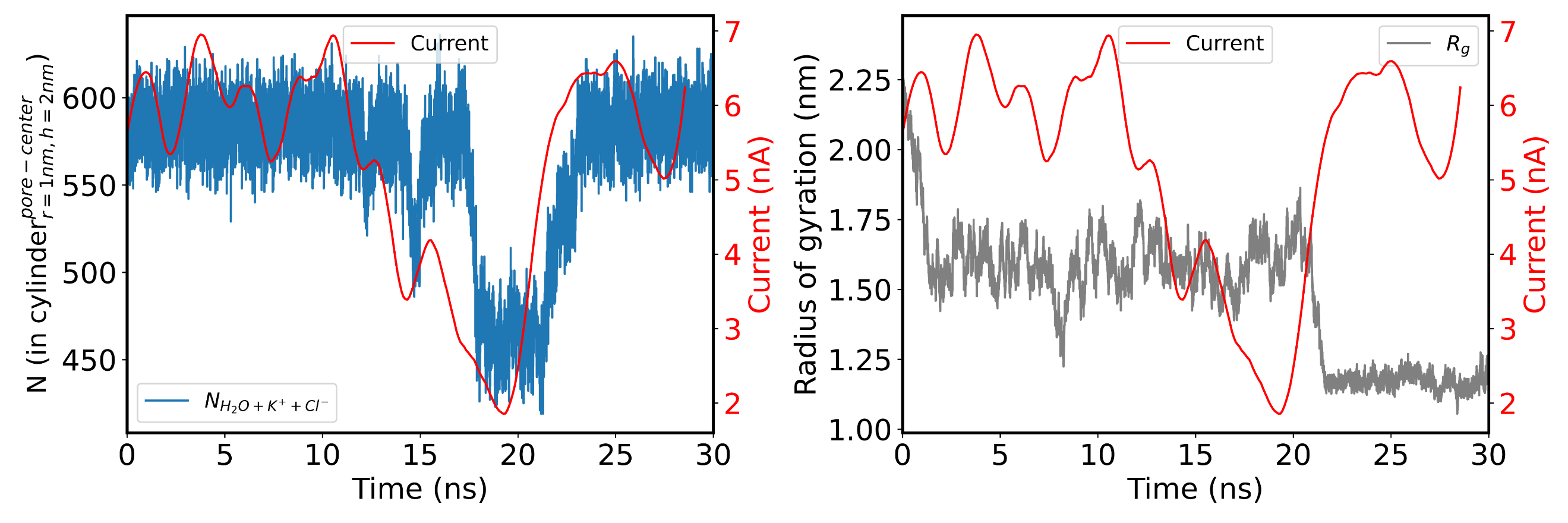

**(b)**

**(a)**

**Fig.S2**: Processed current signal (red curve) exhibiting the dip during translocation. The dip correlates with the reduction of no. of waters and ions (Fig.S2a) inside a cylinder of 2nm height and 1nm radius about the pore, while translocation is happening.

The noisy current signal in Fig.S1 was processed by block averaging the raw data over 100 ps window followed by a moving average of 1.5 ns window and smoothening with Sav-Gol filter which resulted in the red curve shown in **Fig.S2**. As the peptide translocates through the pore, the number of waters and ions inside the pore decreases as well as the value of current (**Fig.S2 a**). After the completion of translocation, the linear peptides tend to coil up (**Video S10**) in quite a few cases and hence, the decrease in radius of gyration (**Fig.S2 b**)

. **2. Assessing underlying distribution for dwell time, relative current blockade and goodness of fit**

In **Fig.3**, we fitted the dwell time with log-normal distribution and relative current blockade with normal distribution. In the following table, we provide a statistical rationale for doing so.

**Dwell time**

Standard Kolmogorov-Smirnov test^[1]^ yields p>0.05 in all cases of log-normal fit and p>0.05 in all cases of normal fit except Ac_apart. Although the normal fit fails to reject the null hypothesis, for six cases, NoPTM, Met barely passes the criteria. To measure the relative goodness of fit among normal and log-normal distribution, we calculate the Akaike Information Criterion^[2]^ (AIC). Out of 7 cases, 5 yield a lower AIC for log-normal distribution which indicates that log-normal is a better fit compared to normal. The two cases where AIC for normal fit was lower, the difference was small. Hence, we chose log-normal fit for dwell time distribution.

**Table S4**

| **PTM** | **Dwell Time** | | | | | | **Relative Current Blockade** | | | | | |
| --- | --- | --- | --- | --- | --- | --- | --- | --- | --- | --- | --- | --- |
|  | **Log-normal** | | | **Normal** | | | **Log-Normal** | | | **Normal** | | |
|  | **KS Statistic** | **p-value** | **AIC** | **KS Statistic** | **p-value** | **AIC** | **KS Statistic** | **p-value** | **AIC** | **KS Statistic** | **p-value** | **AIC** |
| NoPTM | 0.05 | 0.93 | **384.33** | 0.13 | 0.09 | 414.56 | 0.04 | 0.98 | **-272.29** | 0.05 | 0.92 | -271.6 |
| Ac | 0.01 | 0.91 | **222.34** | 0.18 | 0.20 | 237.45 | 0.11 | 0.80 | -76.49 | 0.13 | 0.65 | **-76.77** |
| Ac_apart | 0.11 | 0.83 | **239.05** | 0.25 | 0.04 | 262.85 | 0.06 | 1.00 | -65.88 | 0.08 | 0.99 | **-66.86** |
| Ac_adjacent | 0.08 | 0.78 | 465.49 | 0.06 | 0.96 | **463.28** | 0.08 | 0.78 | -154.64 | 0.1 | 0.52 | **-155.78** |
| Ph | 0.1 | 0.33 | 549.15 | 0.09 | 0.49 | **546.07** | 0.07 | 0.77 | -172.42 | 0.09 | 0.48 | **-172.649** |
| Met | 0.16 | 0.49 | **116.19** | 0.26 | 0.06 | 135.36 | 0.12 | 0.82 | -72.58 | 0.11 | 0.92 | **-76.1** |
| Met_double | 0.19 | 0.82 | **55.36** | 0.31 | 0.25 | 62.12 | 0.15 | **0.95** | -39.81 | 0.14 | 0.97 | **-41.8** |

**Relative Current Blockade**

Standard Kolmogorov-Smirnov test yields p>0.05 in all cases of normal and log-normal fit. The AIC values subjected to normal fit are slightly lower compared to log-normal fit for six out of seven cases. Hence, we chose a normal fit for the relative current blockade distribution.

**3. Indistinguishability among unmodified and methylated peptides**

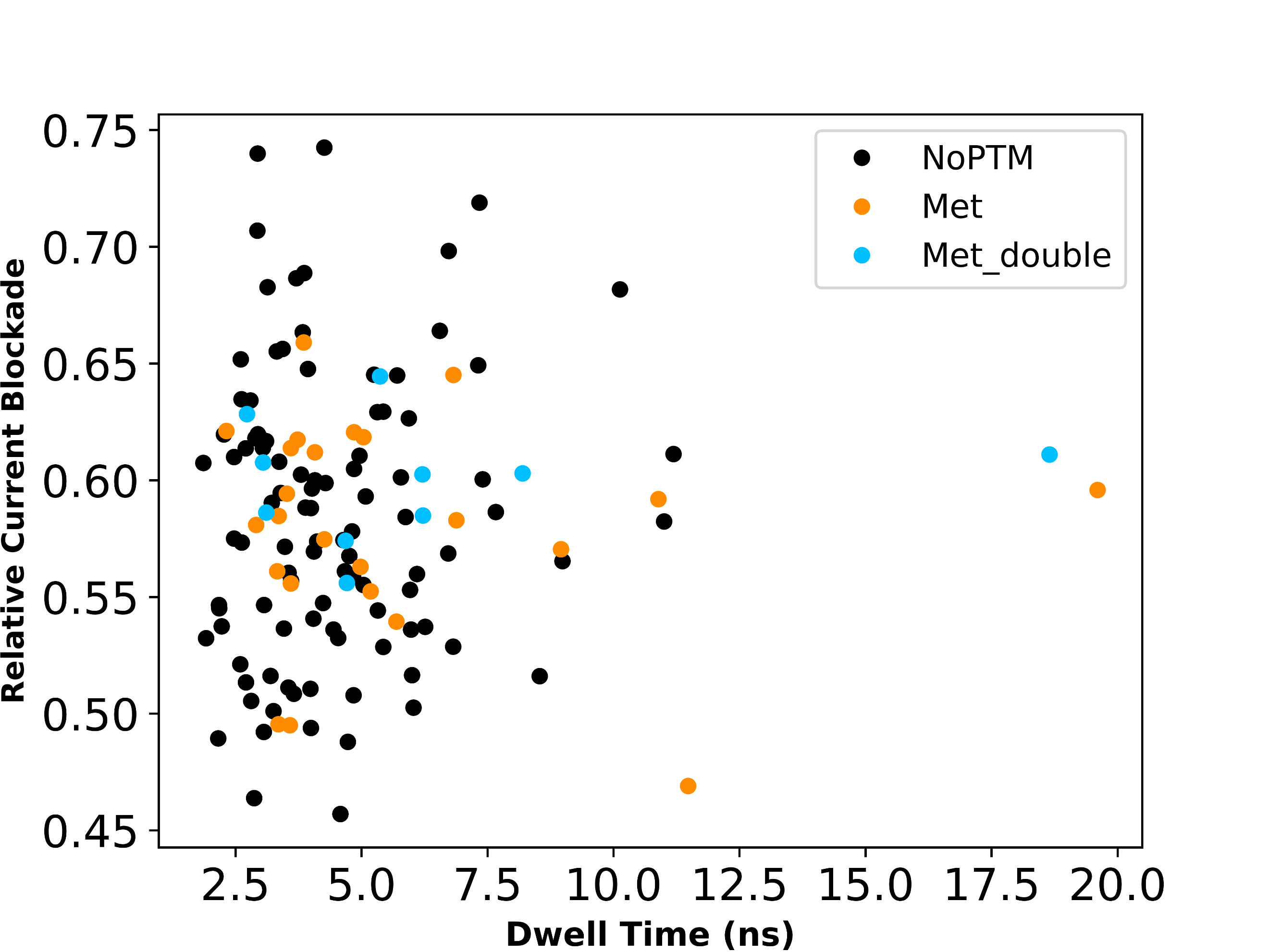

**Fig.S3**: No distinct cluster for NoPTM, Met, Met_double in the dwell time-relative current blockade space.

**4. Supplementary Video**

1. **Video S1**: Translocation of a phosphorylated peptide (Ph). An electrostatic tug-of-war can be observed around pore during translocation.

2. **Video S2**: Translocation of a methylated peptide (Met). Minimal pore-interaction of the methylated residue is observed.

3. **Video S3**: Translocation of a doubly methylated peptide (Met_double). Minimal pore-adherence of the methylated residues is observed.

4. **Video S4**: Translocation of an acetylated peptide (Ac). Enhanced pore-interaction of the acetylated residue is noticeable.

5. **Video S5**: Translocation of a doubly and adjacently acetylated peptide (Ac_adjacent). Enhanced pore-adherence subjected to one of the acetylated residues dominantly resists peptide motion.

6. **Video S6**: Translocation of a doubly and adjacently acetylated peptide (Ac_adjacent). Both acetylated residues engage in strong, synchronous interactions with the pore.

7. **Video S7**: Translocation of a doubly and non-adjacently acetylated peptide (Ac_apart). The second acetylated residue to approach the pore shows longer pore adherence compared to the first.

8. **Video S8**: Translocation of a doubly and non-adjacently acetylated peptide (Ac_apart). Both acetylated residues engage in strong, synchronous interactions with the pore.

9. **Video S9**: Translocation of a doubly and non-adjacently acetylated peptide (Ac_apart). The first acetylated residue to approach the pore adheres to the graphene surface. The next one does not.

10. **Video S10**: Translocation of a unmodified peptide (NoPTM). Minimal pore-interaction is noticeable.
